# Quantum energy levels of glutamate modulate neural biophotonic signals

**DOI:** 10.1101/505024

**Authors:** Zhengrong Han, Weitai Chai, Zhuo Wang, Fangyan Xiao, Jiapei Dai

## Abstract

Glutamate is the most abundant excitatory neurotransmitter in the brain, and it plays an essential and important role in neural functions. Hypofunction of the glutamatergic pathway and the changes in the glutamate-glutamine cycle function are important neuropathological mechanisms of severe mental disorders including schizophrenia and depression. Current studies have shown that glutamate can induce neural biophotonic activity and transmission, which may involve the mechanism of photon quantum brain; however, it is unclear whether such a mechanism follows the principle of quantum mechanics. Here we show that the action of glutamate on its receptors leads to a decrease in its quantum energy levels, and glutamate then partially or completely loses its function to further induce the biophotonic activity in mouse brain slices. The reduced quantum energy levels of glutamate can be restored by direct-current electrical discharges and the use of energy transfer of chloroplast photosynthesis; hence, the quantum energy recovered glutamate can again induce significant biophotonic activity. Furthermore, the changes in quantum energy levels of glutamate are related to the exchange and transfer of electron energy on its active hydrogen atom. These findings suggest that the glutamate-induced neural biophotonic signals may be involved in the transfer of the quantum energy levels of glutamate, which implies a quantum mechanism of neurotransmitter action. The process of glutamate recycling that is related to the synergism of neurons and glial cells and certain key enzymes may be necessary for the recovery of quantum energy levels of glutamate after completion of the neural signal transmission. These findings may also provide a new idea to develop “quantum drugs”.

## Introduction

Analysis of the binding characteristics of biological signaling molecules to their targets or receptors provides an important means to understand their physiological roles and to develop target drugs. However, a large number of studies have found that the effects of signaling molecules on their targets are often unpredictable, for example, the signal molecules or drugs with identical or similar structures such as chiral drugs may produce different effects, while the molecules with completely different structures may induce identical or similar effects (*1*). For olfactory molecules, human beings can directly experience their differences, and quite different molecules can have similar odors, whereas similar molecules can have dissimilar odors (*2*).

The theoretical studies speculate that the reasons for these phenomena may be related to the mechanism of quantum biological action (*3-5*), however, the theoretical framework based on traditional quantum mechanics, such as quantum coherence, entanglement and superposition, could not explain the existence of phenomena very well (*6, 7*). Recent studies have shown that biophoton activity may play an important role in quantum biological mechanisms, and it was reported that glutamate, the most abundant excitatory neurotransmitter in the brain, could induce biophotonic activities and transmission in neural circuits, which may involve the mechanism of photon quantum brain (*8-12*). The characteristic differences such as spectral redshift in glutamate-induced biophotonic activity in different animals may give some evolutionary advantages such as human intelligence (*13*), but may also allow human beings to be more sensitive to the development of neurological and psychiatric diseases due to the changes in neurotransmitter functions. For example, hypofunction of the glutamatergic pathway and the changes in the glutamate-glutamine cycle function are important neuropathological mechanisms of severe mental disorders including schizophrenia and depression (*14-21*). Therefore, we hypothesize that the enhancement or decrease of glutamate function may be related to the changes in quantum energy levels of glutamate, which were experimentally investigated in this study.

## Results

### NMDA receptor mediates glutamate-induced biophotonic activity

Our previous study demonstrated that 50 mM purchased glutamate (p-Glu) can lead to biophotonic activity and transmission (neural biophotonic signals) in neural circuits, presenting four characteristic stages (initiation, maintenance, washing and reapplication) (*8*). In this study, we further found that N-Methyl-D-aspartate (NMDA), a specific agonist at one of the ionotropic glutamate receptors, could also induce similar patterns of biophotonic activity in mouse brain slices (Fig. 1A), suggesting that the NMDA receptor mediates glutamate-induced biophotonic activity. In addition, the washing effect, which provides a further increase in biophotonic activity, was significantly inhibited by the application of a presynaptic vesicle release inhibitor (H89) at the beginning of washing (Fig. 1B and Table S1), suggesting that the washing effect is due to the enhanced presynaptic glutamate release. A possible mechanism is that the extracellular application of glutamate can reduce the release of presynaptic vesicles through an inhibitory effect on the presynaptic membrane (*22*), which results in increased glutamate storage in the presynaptic vesicles and, therefore, a rapid and massive glutamate release from the presynaptic membrane occurs during the early stage of washing.

**Fig. 1.**
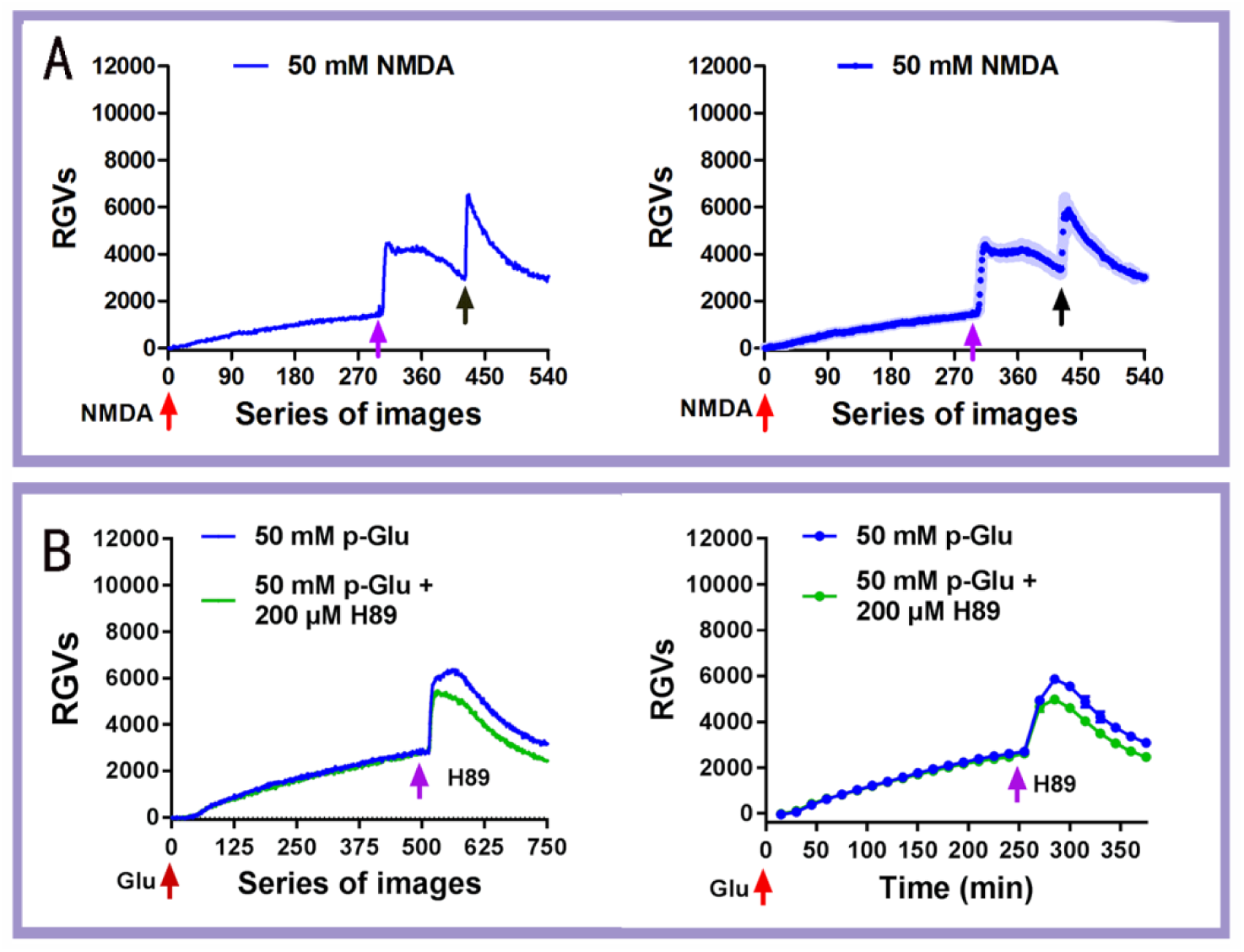
NMDA receptor mediates glutamate-induced biophotonic activity. (**A**) Representative dynamic changes in biophotonic activity demonstrated by the relative grey values (RGVs) in the left panel after the application of 50 mM NMDA, showing four characteristic stages (initiation, maintenance, washing and reapplication; 30 s imaging time for each original image). The population average dynamic changes in biophotonic activity demonstrated by RGVs are in the right panel (n=5). Shaded regions indicate s.e.m. values. The red, pink and black arrows indicate the time points for the application (0 min), washing (150 min) and reapplication (210 min) of NMDA, respectively. (**B**) Representative dynamic changes in biophotonic activity demonstrated by the RGVs in the left panel, showing three characteristic stages (initiation, maintenance and washing; 30 s imaging time for each original image). The right panel shows the sum of the time course of the average change in RGVs from 15 continuously processed original grey images. The red and pink arrows indicate the time points for the application of glutamate (0 min) and 200 μM H89 (240 min) just before washing. The intensities of biophotonic activity tended to decrease significantly after the application of H89 compared to those in controls (n=6 for 50 mM p-Glu and 50 mM p-Glu+200 μM H89). Data are shown as the mean±s. e. m. n=the number of slices from the same number of mice. Significant differences (** p<0.01) are noted from 271-285 min (see Table S1).

### Decrease in the quantum energy levels of glutamate after its action

Then, we observed long-term (24 h) effects of 50 mM glutamate-induced biophotonic activity. We designed a consumption experiment to evaluate the changing patterns of glutamate-induced biophotonic activity. Mouse brain coronal slices (450 µm in thickness) were cut from the anterior part of the olfactory cortex to the caudal part of cerebellar cortex and all slices were collected. We found that the intensities of glutamate-induced biophotonic activities from a slice for imaging, which was perfused with 100 ml ACSF containing 50 mM glutamate together with four whole mouse brain slices in a bottle, dramatically decreased after perfusion and imaging for approximately 12 h (12.02±0.65 h, Fig. S1). This result allowed us to speculate that the cause of the obvious weakening of glutamate action may be related to the decrease in the quantum energy levels of glutamate after its action on the postsynaptic receptors (Fig. 2A) because a cyclic perfusion with a fixed volume of ACSF containing 50 mM glutamate was able to partially and almost completely exhaust the action of glutamate by providing appropriate brain tissues and enough perfusion time.

**Fig. 2.**
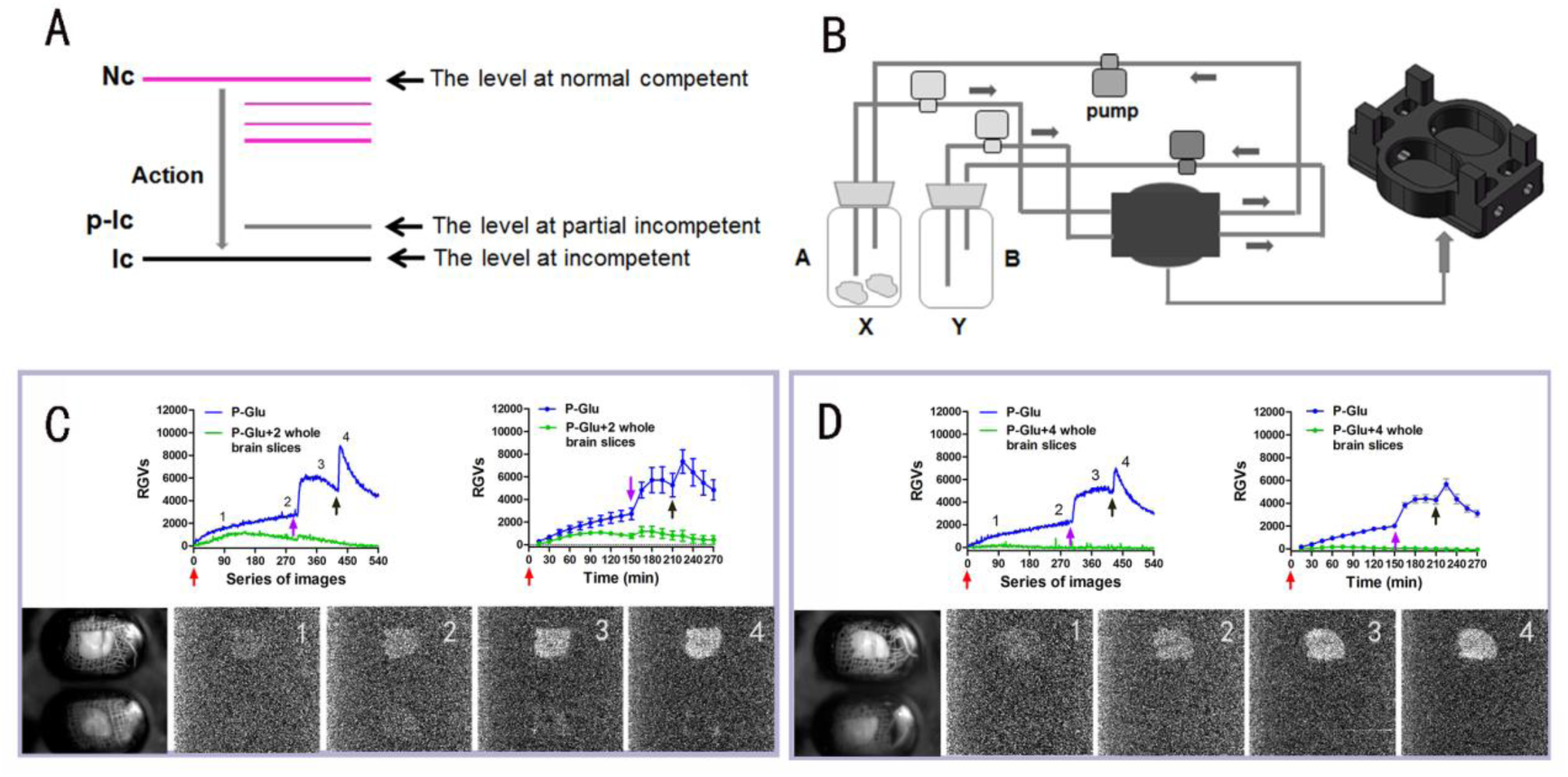
Loss of the glutamate effect is due to the decrease in quantum energy levels. (**A**)Schematic drawing of the decrease in quantum energy levels from the level at the normal competent (Nc) to the levels of partial and complete loss of quantum energy after the glutamate action on its receptors. Ic: incompetent; p-Ic: partial incompetent. (**B**) Schematic drawing of cyclic dual perfusion system, named perfusion system **A** and **B**. The perfusion chamber under the same imaging field is separated into two smaller perfusion parts (subchambers) (see the figure, the right part of the panel), allowing support of independent cyclic perfusion for the left and right hemisphere slices together with two perfusion solution storage glass bottles (**X** and **Y**) and input and output micropumps. **(C, D)** Representative dynamic changes in biophotonic activity demonstrated by the RGVs are shown in the upper left panel. The sum of the time course of the average change in RGVs from 30 continuously processed original grey images is shown in the upper right panel. One of the two hemisphere slices (lower slice in the lower left regular photograph) was perfused independently with 50 ml of ACSF containing 50 mM p-Glu after treatment with 2 whole brain slices (**C**) or 4 whole brain slices (**D**) for 24 h. Another one (upper slice in the lower left regular photograph) was also perfused independently with 50 ml of ACSF containing 50 mM p-Glu as a control. Representative biophoton grey images at the selected time periods indicated in the lower panel and digit: 1-4 mark the 90, 270, 360 and 450 series of images, respectively (30 s imaging time for each image). The red, pink and black arrows indicate the time points for application of perfusion solution (0 min), washing (150 min) and reapplication (210 min), respectively. The intensities of biophotonic activity were decreased significantly after treatment with 2 whole brain slices for 24 h (green lines) compared to those of the controls (blue lines, **c**, n=6), and were almost completely eliminated with 4 whole brain slices (**D**, n=6). p-Glu: purchased glutamate. Data are shown as the mean±s. e. m. n=the number of slices from the same number of mice. Significant differences (* p<0.05 or ** p<0.01) are noted from 91-105 min for **C** and 1-15 min for **D** (see also Table S1).

To confirm our results, we designed a cyclic dual perfusion system to conduct further experiments. Two hemisphere slices (left and right, 450 µm in thickness) prepared from a coronal whole brain slice at the anterior level of the hippocampus were placed separately into two perfusion subchambers that were imaged simultaneously (Fig. 2B). We found that the patterns and intensities of 50 mM glutamate-induced biophotonic activity were exactly the same for both slices (Fig. S2), which was consistent with data obtained from a whole brain coronal slice containing left and right hemispheres (*8*). Then, we tested the effects by only using ACSF containing 50 mM p-Glu after incubation with different quantities of brain slices. All slices prepared from two or four mouse brains were incubated in 100 ml ACSF containing 50 mM p-Glu for 24 h at ~ 28℃. Then, 50 ml incubated solution was collected to perfuse a new prepared brain slice for biophotonic imaging. We found that the intensity of biophotonic activity during the four stages induced by the solution after incubation with two whole brain slices was significantly weaker than that in control solution (Fig. 2C and Table S1), while the effects were almost eliminated after incubation with four whole brain slices, both of which resulted in hypofunctional (partial incompetent, p-Ic) and nonfunctional glutamate or incompetent glutamate (Ic-Glu) (Fig. 2D and Table S1). The observed results were not due to the change in the molecular structure of glutamate after the action of its receptors or the effects of other active molecules released from the brain slices during incubation. Even if the low concentration of classical neurotransmitters such as acetylcholine, dopamine, 5-hydroxytryptamine (5-HT) and norepinephrine were present in the ACSF after incubation for 24 h, their presence cannot explain the weakening and disappearance of glutamate action because our previous study demonstrated that even high concentrations of 5-HT (> 20 µM) only produced partial inhibitory effects on 50 mM glutamate-induced biophotonic activity, while acetylcholine, dopamine and norepinephrine played reinforcing rather than inhibitory roles (*12*). Therefore, in essence, these results proved our speculation mentioned above.

### Electrical discharge restores the quantum energy levels and the effects of incompetent glutamate

To test whether there are ways to restore the decreased quantum energy levels of glutamate and reverse its action and function to induce biophotonic activity, we developed two techniques (Fig. 3A). One is called the electrical stimulation assisted quantum energy enhancement technique (ESA-QEET, Fig. 3B, see also SI for details). One hundred millilitres ACSF containing 50 mM p-Glu was first incubated with four whole brain slices for 24 h at ~ 28°C and then divided into two parts (50 ml each). One portion of the solution was treated for 3 h with 12 V direct-current electrical discharges (DC-ED) at a high frequency (~10 k Hz), and another was the control. We found that such treatments significantly restored the biophotonic activity of Ic-Glu compared to the controls without DC-ED treatment and reached to 59.00-118.25% of the effects of ACSF containing 50 mM p-Glu at the different imaging periods (Fig. 3C and Table S1). The recovery of the quantum energy level of glutamate was not due to the influence of DC-ED on the other components of the ACSF because the ACSF without p-Glu incubated with four whole brain slices for 24 h did not have any effects on the biophotonic activity after treatment with 12 V DC-ED (Fig. 3D and Table S1). These findings also imply that the increase in quantum energy levels of glutamate may be involved in the transfer of electron energy on its molecule, and a possible explanation is that the active hydrogen (proton) of glutamate is responsible for the exchange and transfer of quantum energy, which was supported by our findings that, similar to 50 mM p-Glu, 50 mM acetic acid (0.3%) could also induce similar patterns of biophotonic activity in mouse brain slices (Fig. S3 and Table S1), whereas 50 mM sodium glutamate (gourmet powder), on which the sodium atom is substituted for the active hydrogen atom, could not have any effect on the induction of biophotonic activity in mouse brain slices (Fig. S4 and Table S1).

**Fig. 3.**
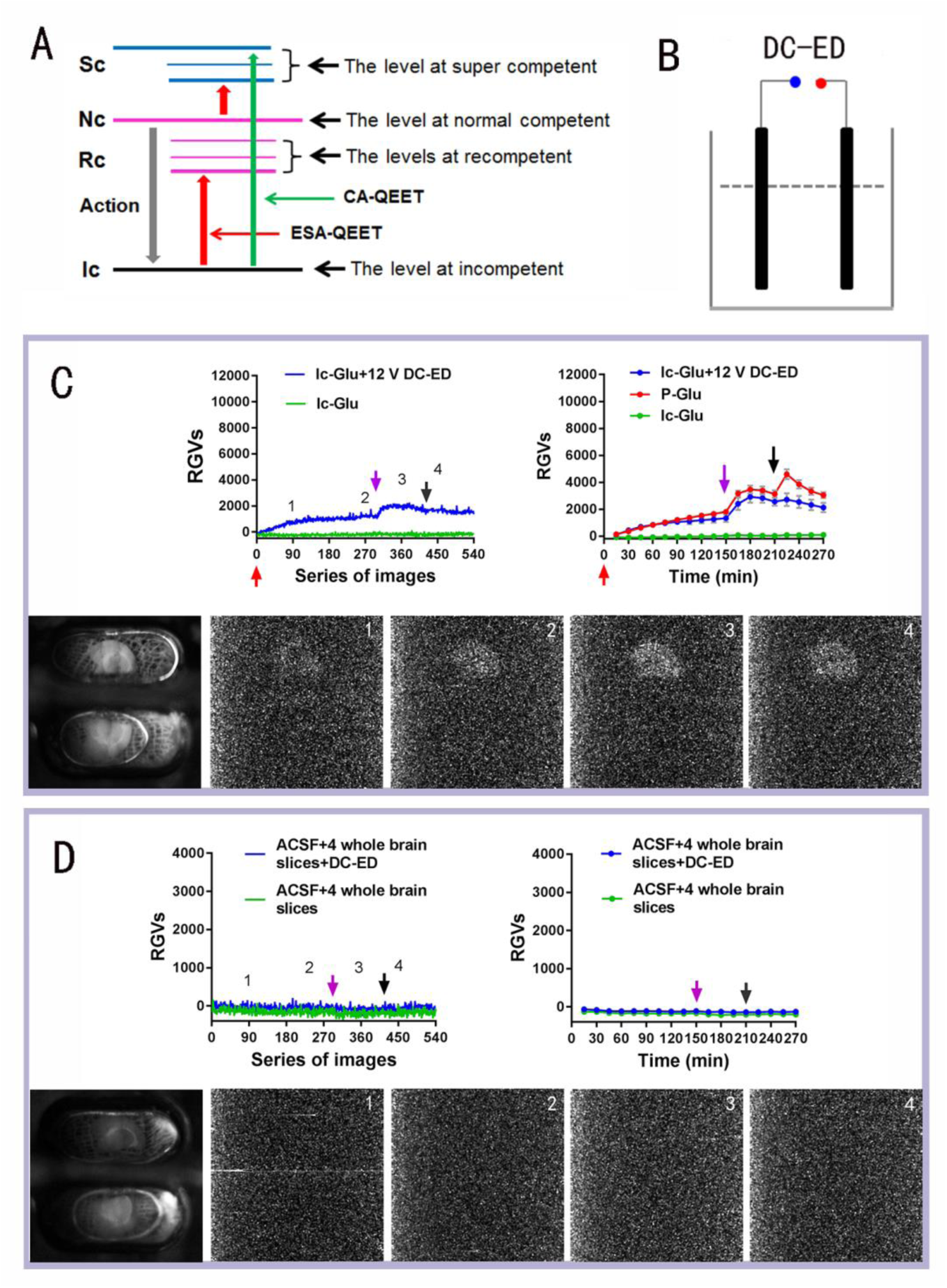
Electrical discharge restores the quantum energy levels and the effects of incompetent glutamate. (**A**) Schematic drawing of the potential recovery of the quantum energy levels (Rc) of glutamate after treatment with the electrical stimulation assisted quantum energy enhancement technique (ESA-QEET, lower red arrow) or chloroplast-assisted quantum energy enhancement technique (CA-QEET). The levels at super competent (Sc) can also be obtained by ESA-QEET from the normal level (upper red arrow) or by CA-QEET from the incompetent level (blue arrow). (**B**) Schematic drawing of the ESA-QEET. A power supply device with 12 V direct-current electrical discharges (DC-ED) at a high frequency (~10 k Hz) was used to execute the electrical stimulations using the two same carbon rods as the cathode and anode. **(C, D)** Representative dynamic changes in the biophotonic activity demonstrated by the RGVs are shown in the upper left panel. The sum of the time course of the average change in RGVs from 30 continuously processed original grey images is shown in the upper right panel. One of two hemisphere slices (upper slice in the lower left regular photograph) was perfused independently with the recompetent glutamate treated by 12 V DC-ED for 3 h after its incompetent treatment with 4 whole brain slices for 24 h (**C**) or ACSF treated by 12 V DC-ED for 3 h after incubation with 4 whole mouse brain slices for 24 h (**D**). Another hemisphere slice (lower slice in the lower left regular photograph) was perfused independently with the incompetent glutamate without DC-ED treatment (**C**) or ACSF without treatment by 12 V DC-ED after incubation with 4 whole mouse brain slices for 24 h (**D**) and considered the controls. Representative biophoton grey images at the selected time periods indicated in the upper left panel and digit: 1-4 mark the 90, 270, 360 and 450 series of images, respectively (30 s imaging time for each image). The red, pink and black arrows indicate the time points for application of the perfusion solution (0 min), washing (150 min) and reapplication (210 min), respectively. Approximately 59.00-118.25% of the effects of p-Glu (red line) were reached at the different time imaging periods after treatment for 3 h with 12 V DC-ED (n=8) compared to the reference induced by 50 mM p-Glu during the different imaging periods (n=37) (**C**). No significant increase in biophotonic activity in four stages after treatment with DC-ED (n=5) was observed compared to that without treatment (**D**). p-Glu: purchased glutamate. Data are shown as the mean±s. e. m. n= the number of slices from the same number of mice. Significant differences (* p<0.05 or ** p<0.01) are noted from 1-15 min for c (see also Table S1).

### Chloroplast photosynthesis restores the quantum energy levels and the effects of incompetent glutamate

Another technique is called the chloroplast-assisted quantum energy enhancement technique (CA-QEET, Fig. 3A, see also SI for details). Chloroplasts can achieve efficient light energy transfer through photosynthesis under light illumination and, therefore, the electron and proton transport between molecules can be realized, in which quantum mechanics is involved (*23-26*). We explored whether chloroplasts were able to allow the hypofunctional or nonfunctional glutamate to restore its quantum energy levels. Chloroplasts were prepared from a vegetable (spinach) and a comparative analysis showed that the isolated chloroplasts could maintain a certain degree of light transfer activity in the ACSF and the ACSF containing 50 mM p-Glu compared to those under an appropriate condition reported previously (*27*) (Fig. S5). We added ~3 g freshly prepared chloroplast precipitate into 50 ml ACSF containing Ic-Glu in two steps (~1.5 g each) and incubated the mixed preparations for 48 h under a combination of day and night illumination (see SI for details). We found that such treatments allowed Ic-Glu to reach 160.25-370.70% of the effects of the 50 mM p-Glu at the different imaging periods (Fig. 4A and Table S2), showing the super recompetent effects (Fig. 3A). The over-recovery of biophotonic activity was partially because of the acidification effect of light energy transfer of the chloroplasts on ACSF (*28-30*), which resulted in low pH (from ~7.4 to ~4.0) and induced some degree of biophotonic activity both from the ACSF treated with and without four whole brain slices (Fig. 4B and Table S2); however, adjusting the acidified ACSF to the original level of pH 7.4 significantly reduced the effects and only induced slight biophotonic activity (Fig. 4C and Table S2), reinforcing the role of the active hydrogen for the glutamate-induced biophotonic activity mentioned above.

**Fig. 4.**
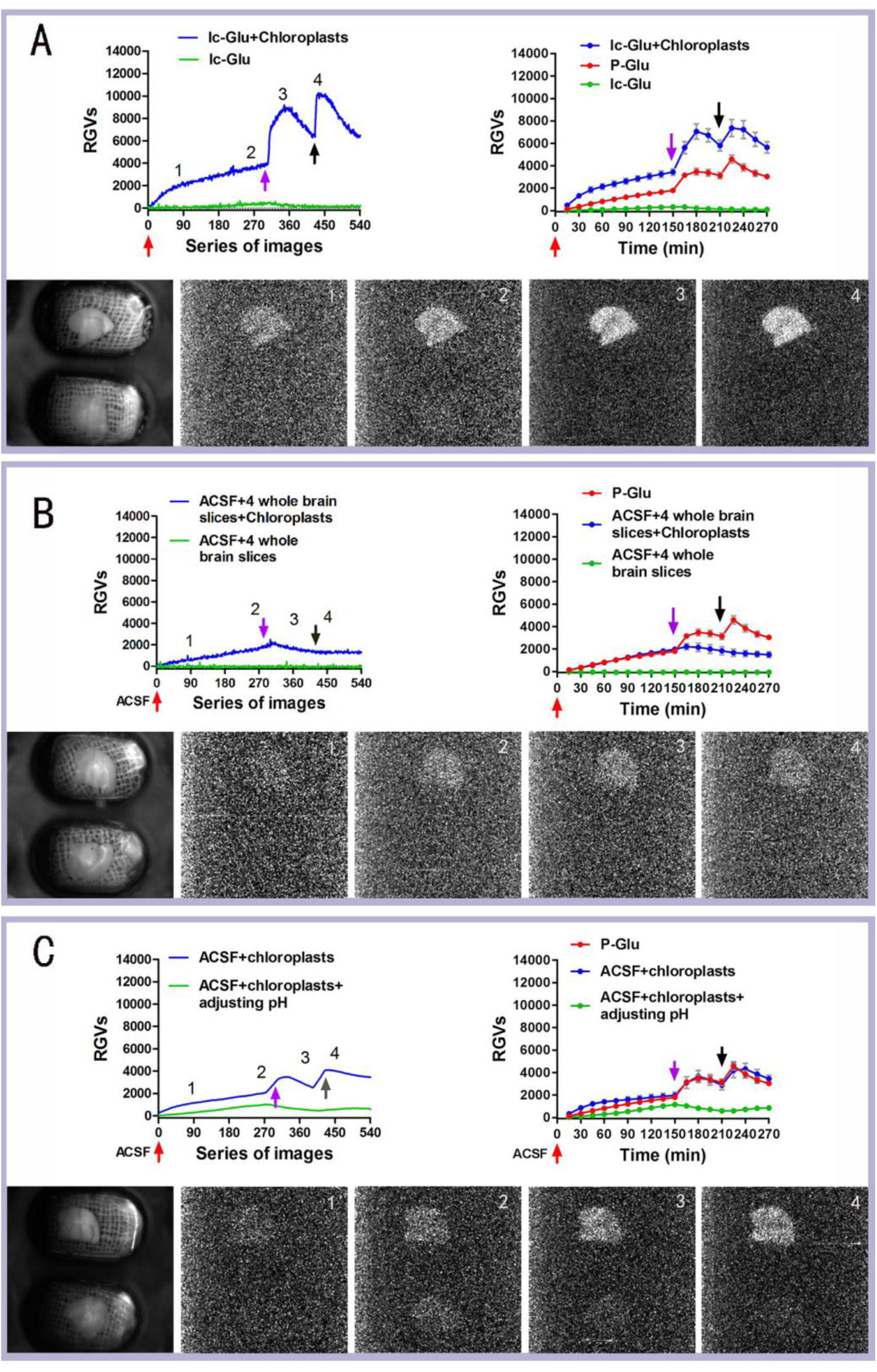
Chloroplast photosynthesis restores the quantum energy levels and the effects of incompetent glutamate. **(A-C)** Representative dynamic changes in biophotonic activity demonstrated by the RGVs are shown in the upper left panel. The sum of the time course of the average change in the RGVs from 30 continuously processed original grey images is shown in the upper right panel. One of two hemisphere slices (upper slice in the lower left regular photograph) was perfused independently with the recompetent glutamate treated with ~3 g freshly prepared chloroplasts for 48 h under light illumination after glutamate’s incompetent treatment with 4 whole brain slices for 24 h. Another slice (lower slice in the lower left regular photograph) was perfused independently with the incompetent treatment without chloroplast incubation. Representative biophoton grey images at the selected time periods indicated in the upper left panel and digit: 1-4 mark the 90, 270, 360, and 450 series of images, respectively (30 s imaging time for each image). The red, pink and black arrows indicate the time points for application of perfusion solution (0 min), washing (150 min) and reapplication (210 min), respectively. (**A**) A 160.25-370.70% increase in biophotonic activity was obtained for incompetent glutamate (n=5) compared to the reference induced by 50 mM p-Glu at the different imaging periods (n=37). (**B**), The biophotonic activity was increased for the ACSF treatment with ~3 g of freshly prepared chloroplasts after incubation with 4 whole brain slices for 24 h but were less than that in **A** (n=5). (**C**) ACSF treated with ~3 g of freshly prepared chloroplasts for 48 h resulted in increased biophotonic activity (n=5), which was reversed by adjusting the acidified ACSF to the original level of pH 7.4, indicating the acidification of ACSF by chloroplast. Data are shown as the mean±s. e. m. n= the number of slices from the same number of mice. The reference data from 50 mM p-Glu in **A** (right panel) are the same samples as in Fig. 3C (right panel). Significant differences (*p<0.05 or ** p<0.01) are noted for **A-C** (see also Table S2).

## Discussion

This study demonstrated that, after induction of neural biophotonic signals, glutamate lost its action and presented a decline in the molecular quantum energy levels, but it did not involve the changes in its molecular structure. Although the exact quantum biological mechanism, including the atoms involved, is unclear, our findings suggest that this phenomenon may be highly associated with the active hydrogen atom on the glutamate molecule. The change in the quantum states (quantum levels) in a liquid molecule involves the translational level of the molecule, the rotation, vibration and motion of the electrons, and the change in the energy level of the nuclear motion in the molecule (*31-33*). The mechanism of the decrease in the quantum energy levels after the action of glutamate on its receptors may be mainly related to the change in the electron quantum energy levels in the active hydrogen atom.

Therefore, our findings may provide a new idea for further clarifying the importance of the quantum biological mechanism of the functional molecules in the brain. For example, the glutamate released into the synaptic cleft is reused mostly by uptake into surrounding glial cells via the glutamate-glutamine (Glu-Gln) cycle, which is related to the synergism of neurons and glial cells, glutamate transporter and certain key enzymes (*34, 35*), and this widely-known process of recycling of glutamate seems to violate the energy conservation principle of the organism. Our findings suggest that such a process of recycling is indeed a necessary step for recovering the quantum energy levels of glutamate after its action, possibly as an important mechanism of quantum biology. Second, our findings could lead to a new perspective for elucidating the pathophysiological mechanisms of neuropsychiatric disorders. The changes in the glutamate recirculation, including the changes in transporters and key enzymes, are related to the pathological mechanisms of these diseases (*14-21*), while the action of an enzyme is considered to be a quantum mechanism, which is involved in the transfer of quantum energy (*3*). Therefore, the disruption of the Glu-Gln cycle may play an important role in the hypofunction of the glutamatergic pathway and contribute to the pathological mechanisms of neuropsychiatric disorders. Finally, our findings could provide new explanations for some clinical therapies such as local brain electromagnetic stimulations, electroshock, traditional natural medicine and acupuncture treatments (*36, 37*). One of the possible mechanisms for these treatments is through the restoration of the imbalance of the quantum energy levels of neurotransmitters and other functional molecules. In addition, the differences in the quantum energy levels of odorants may provide a novel way of explaining the diversity of the olfactory sense (*2*).

## Methods

The detailed information for materials and methods is described in Supporting Information (SI).

## Acknowledgments

This work was supported by the Sci-Tech Support Plan of Hubei province (2014BEC086) and the research funds of South-Central University for Nationalities (XTZ15014 and CZP 18008), and partially by the National Natural Science Foundation of China (31640034, 31700911).

## Author contributions

J.D. conceived and designed the experiments; R.Z, T.C, and J.D. performed the experiments and analyzed the data; Z.W, and F.X. provided technical support for UBIS and research materials; J.D. wrote manuscript.

## Supporting Information (SI)

### Materials and Methods

#### Experimental mice

Adult male Kunming mice (6-8 weeks), which were originally generated from a Swiss white mouse and has been widely used in biomedical studies in China, were purchased from Hubei Provincial Laboratory Animal Public Service Center (Wuhan, China) and housed in a room with a 12 hrs light/dark cycle (lights on at 7:00AM) with access to food and water adlibitum. The study protocol was approved by the Committee on the Ethics of Experimental Animals and Biomedicine of South-Central University for Nationalities.

#### Preparation of mouse brain slices

Mice were decapitated and the brain was quickly removed and placed in ice-cold (0-4°C) artificial cerebral spinal fluid (ACSF). ACSF contained (in mM) 125 NaCl, 2.5 KCl, 2 CaCl_2_, 1 MgCl_2_, 1.25 NaH_2_PO_4_, 26 NaHCO_3_ and 20 D-glucose; pH 7.4. The concentrations of NaCl and D-glucose were adjusted to 115 mM and 10 mM, respectively, if 50 mM glutamate was added to the ACSF to maintain the relative stability of the osmolality.

The coronal brain slices were cut to 450 µm thickness with a Vibratome (Leica VT1000S, Germany) beginning at the rostral part of the olfactory cortex and ending at the caudal part of the cerebellum. A standardized coronal brain slice or its left and right hemisphere slices at the level of the anterior part of the hippocampus was used for biophoton imaging. The prepared brain slices were kept in ACSF at 0-4°C for further experiments.

#### Cyclic dual perfusion system for brain slices

Schematic drawing of the cyclic dual perfusion system is described in details in Fig. 2b in the main text. The perfusion for two parts of the chamber (subchamber) was maintained independently through an input micropump (~4 ml/min) and an output micropump (6-12 ml/min). A mixture of 95% O_2_+5% CO_2_ was constantly supplied with a membrane oxygenator placed in the ACSF of glass bottle during the perfusion period. The temperature (~35°C) of the medium in the perfusion chamber was maintained with an electrical heater.

#### In vitro biophoton imaging system

The *in vitro* ultra-weak biophoton imaging system (UBIS) was described in our previous studies (*1, 2*). Biophotonic activity (emissions) was detected and imaged with the UBIS using an EMCCD camera (C9100-13 or C9100-23B, Hamamatsu Photonics K. K., Hamamatsu, Japan) in water-cool mode (in this situation, the working temperature at the EMCCD camera can be maintained as low as −95°C) controlled by an image analysis software program (HCImage, Hamamatsu Photonics K. K., Hamamatsu, Japan). The specific steps for biophoton detection and imaging were as follows: (1) the brain slices were transferred to the two subchambers and perfused with ACSF in complete darkness by closing the dark box of the UBIS for approximately 30 min before imaging to exclude the effects of ambient light; (2) real-time imaging was performed by automatically taking an image every 30 s for detection of typical biophotonic activity; (3) the imaging duration was 240-300 min (480-600 series of images) depending on the experimental protocols including an additional 30 min imaging time when the brain slices were only perfused with ACSF before any treatment was carried out; and (4) a regular photograph of brain slices was taken under the normal CCD model before and after the imaging processes were completed to locate its position under the imaging field of view.

#### Image processing, data extraction and analysis

The specific processes for image processing and data analysis were as follows: (1) all original grey images in TIF format were processed with a program running on the MATLAB platform to eliminate the effect of cosmic rays (white spots) as described before (*1*), resulting in the processed biophoton grey images. The parameters for image processing were determined according to the maximum grey values of the biophotonic signals and white spots to eliminate the effect of cosmic rays, but not lose the biophotonic signals; (2) the average grey values (AGVs) of the processed biophoton grey images in the regions of interest (ROIs) were extracted with the same image analysis software program mentioned above, and the data were exported to Microsoft Excel for further analysis. The ROIs included the whole area of the brain slice (slice area) and part of the non-brain slice area (background area); (3) the relative grey values (RGVs) of the brain slice at the different time points were calculated as:

RGVs= AGVs (slice area) - AGV (background area)

The background AGV used for the calculation of RGVs in a given image is chosen from a mean of average grey values from the background ROIs of all processed images.

#### Acquisition of incompetent glutamate

All slices cut from two or four mouse brains were collected and transferred to 100 ml ACSF containing 50 mM glutamate in a glass bottle and incubated for 24 h at ~28 °C. The incubation temperature was maintained with a thermostat water bath. A mixture of 95% O_2_+5% CO_2_ was constantly supplied with a membrane oxygenator placed in the ACSF during the incubation period. The slices were removed from the glass bottle after 24 h incubation and the remaining ACSF was then filtered with a filter paper.

#### Recovery of the quantum energy level of incompetent glutamate by electrical discharges

A direct-current electrical discharge (DC-ED) device with adjustable voltages (Taiyuan Weiyide Automation Technology Co., Ltd., WD-16A) was used to execute electrical discharges (EDs) using the two same carbon rods as the cathode and anode. One hundred millilitres ACSF containing 50 mM glutamate after treatment with four whole brain slices for 24 h was equally divided into two parts (50 ml each). One part was then treated for 3 h with 12 V DC-ED at a high frequency (~10 k Hz) and the other was a control without DC-ED treatment. Same treatments were carried out for one hundred millilitres ACSF without 50 mM glutamate.

#### Isolation of chloroplasts and determination of photochemical activity in the ACSF

Chloroplasts were isolated from spinach according to the methods described previously (*3*). Fresh spinach leaves were obtained from a commercial supplier and homogenized with a homogenizer in a medium containing 50 mM Tris-HCl buffer (pH 7.4), 0.4 mM sucrose and 10 mM NaC1 at room temperature. After filtering through a standardized wire woven mesh test sieve (GB6003.1-1997), the residue was discarded and the filtrate was centrifuged for 3 min at 1000 rpm. The precipitate was removed and the supernatant was centrifuged in the same centrifuge for 5 min at 3000 rpm. The precipitate contained intact chloroplasts and was used for further experiments.

Hill reaction activity and the efficiency of isolated chloroplasts in three reaction solutions (50 ml each) including the reaction mixture [50 mM Tris-HCl buffer (pH 7.4), 0.4 mM sucrose, 10 mM NaC1], ACSF and ACSF containing 50 mM p-Glu were determined by the reduction of 2,6-dichlorophenolindophenol (DCIP). Freshly prepared chloroplast precipitate (1.5 g) was immediately added to each of the three reaction solutions, followed by the addition of 1 ml of 50 mM DCIP (the final reaction concentration of DCIP was ~1 mM), which were then incubated under a combination of day and night illumination (day time: natural window light irradiation from sunrise to sundown, a light intensity of 15,000-50,000 lx; night time from sundown to sunrise: artificially controlled visible light irradiation, a light intensity of ~2000 lx). Every half an hour, the colour changes in the reaction solutions were visually observed and were photographed by a digital camera, with the exception of sleeping hours (from 22:00 PM to 8:00 AM). The same volume (1 ml) of DCIP was added again after approximately 21 h when the colour change from blue to colourless was defined for all three solutions. The whole observation period lasted approximately 28 h. We found that 1.5 g of freshly prepared chloroplast precipitate in 50 ml of ACSF containing 50 mM p-Glu had better efficiency, which remained after incubation for 28 h.

#### Recovery of the quantum energy level of incompetent glutamate and its activation by chloroplast photosynthesis

One hundred millilitres of ACSF containing 50 mM glutamate was incubated with four mouse brain slices for 24 h and then equally divided into two glass bottles (50 ml each). Freshly prepared chloroplast precipitate (1.5 g) was added to one of two glass bottles and incubated for 24 h under a combination of day and night illumination as described above. Then, the chloroplasts were removed from the solution by centrifugation for 5 min at 3000 rpm, and an additional 1.5 g freshly prepared chloroplast precipitate was added and incubated for another 24 h. After centrifuging the chloroplasts, the remaining ACSF supernatant containing recompetent glutamate was tested for biophotonic activity in mouse brain slices as described above.

#### Drug preparation

Glutamate (50 mM, Sigma, St. Louis, MO, USA), NMDA (50 mM, Sigma, St. Louis, MO, USA) and H89 (200 µM, Selleck Chemicals, USA) was directly dissolved in the ACSF. DCIP (50 mM, Shyuanye, Shanghai, China) was initially dissolved in distilled water and then diluted to its final concentration in the incubation solution.

#### Statistical analysis

Statistical analyses were performed using Microsoft Excel, and details of the results are described in Extended Data Table 1-3. Two-tailed Student’s t-test was used to compare the effects at different time points or different time periods in treatment groups.

## Supporting figure legends

**Fig. S1.**
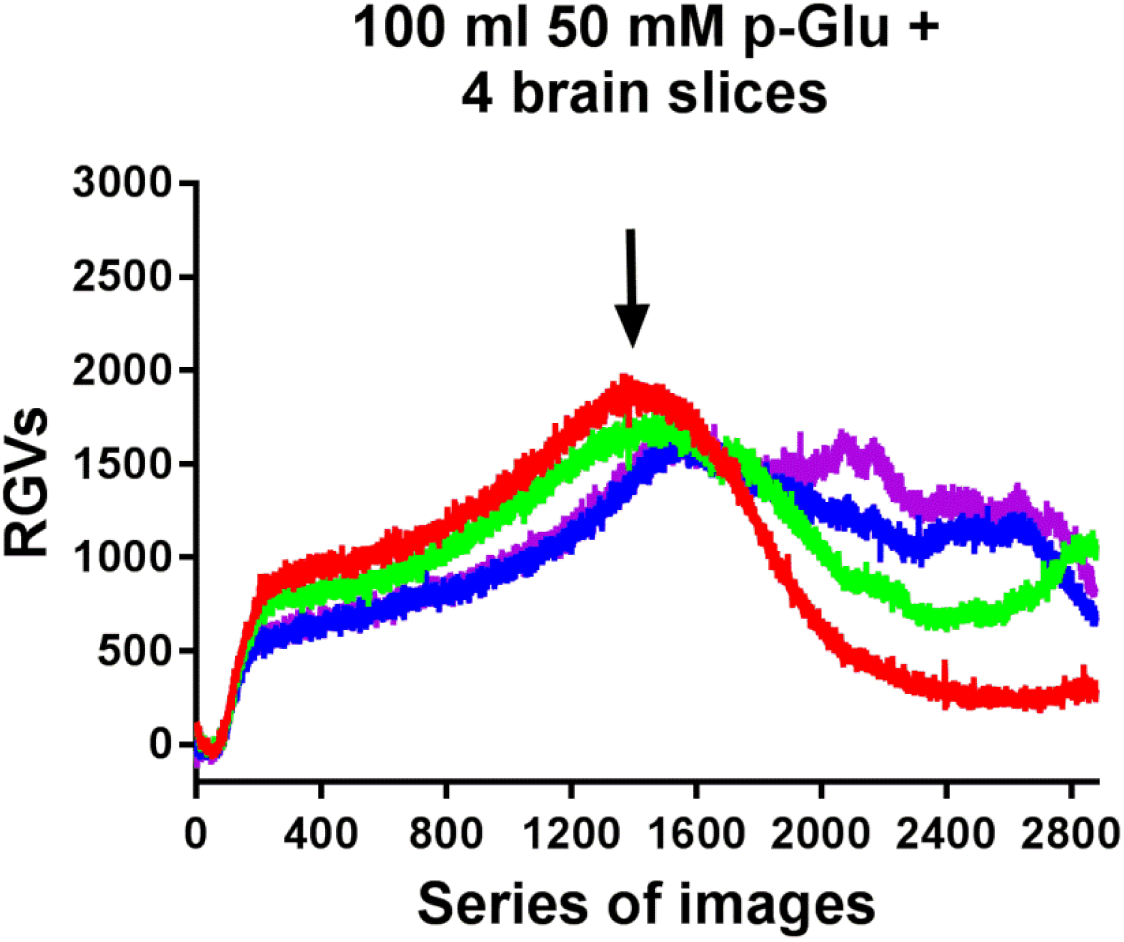
The glutamate-induced biophotonic activity tended to decrease after long-term perfusion with four whole brain slices. Dynamic changes in the biophotonic activity in an imaging slice demonstrated by RGVs are shown after the cyclic perfusion with 100 ml of ACSF containing 50 mM p-Glu and four whole mouse brain slices, which were placed into a perfusion bottle. The intensities of biophotonic activity in four imaging slices (red, green, blue and pink, 30 s imaging time for each image) tended to decrease significantly after approximately 12 h of perfusion (12.02±0.65, n=4). The total imaging time was 24 h for each case.

**Fig. S2.**
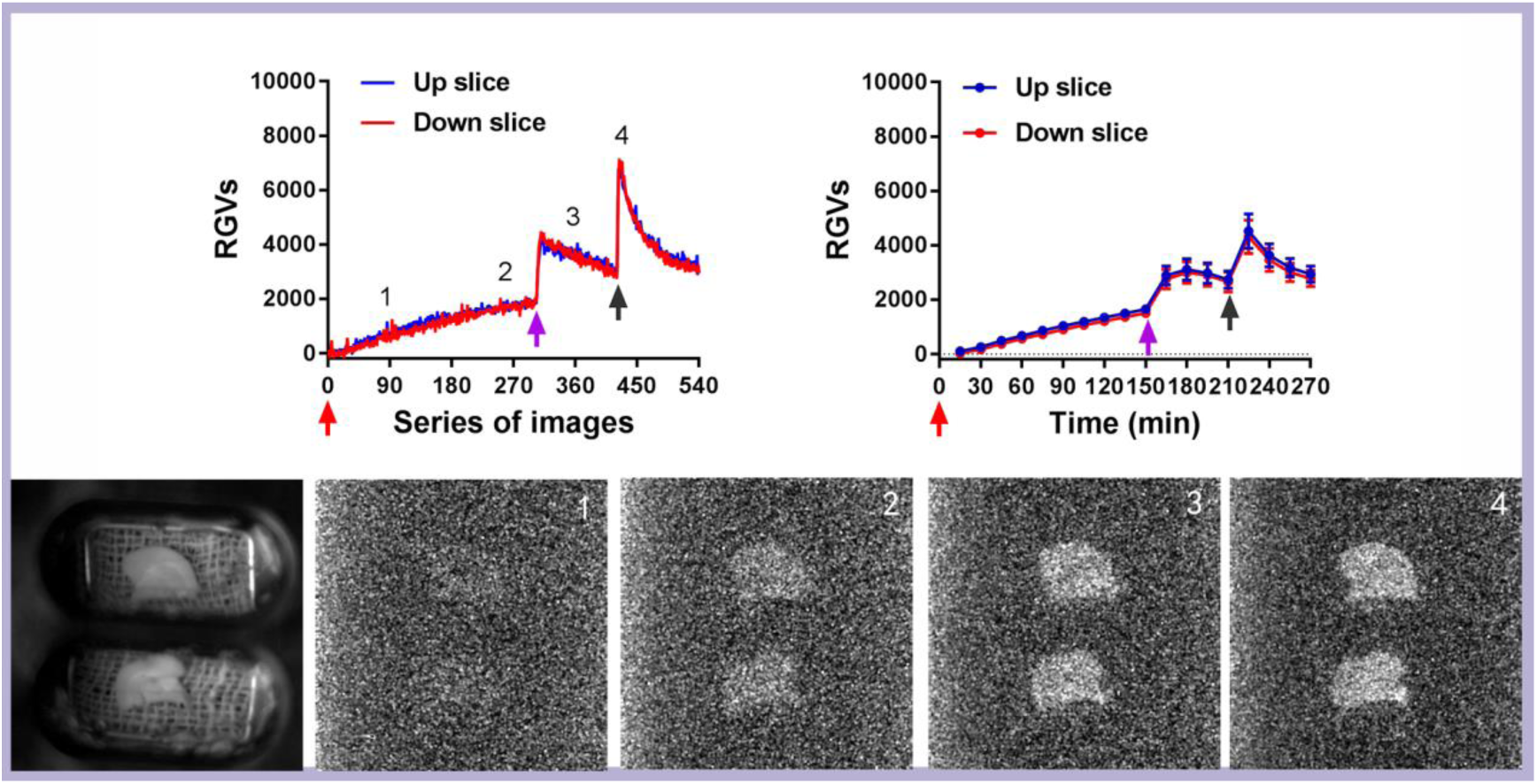
The patterns and intensities of biophotonic activity are the same between the left and right hemisphere slices. Representative dynamic changes in the biophotonic activity demonstrated by RGVs are shown in the upper left panel after the cyclic perfusion of left and right hemisphere slices with 50 ml ACSF containing 50 mM glutamate. The sum of the time course of the average change in RGVs from 30 continuously processed original grey images is shown in the upper right panel (n=10). Red and blue lines indicate the dynamic biophotonic activity in the up and lower slices, respectively. The patterns and intensities of 50 mM glutamate-induced biophotonic activity were the same for both slices, which was consistent with that obtained from a whole brain slice containing left and right hemispheres. Digit: 1-4 mark the 90, 270, 360, and 450 series of images, respectively (30 s imaging time for each image) in the upper left panel, which corresponds to the four images in down panel. The red, pink and black arrows indicate the time points for application of 50 mM glutamate (0 min), washing (150 min) and reapplication (210 min), respectively. A representative regular image of two brain slices (up and down) is shown on the left side of the lower panel. Data are shown as the mean±s. e. m. n=the number of slices from the same number of mice.

**Fig. S3.**
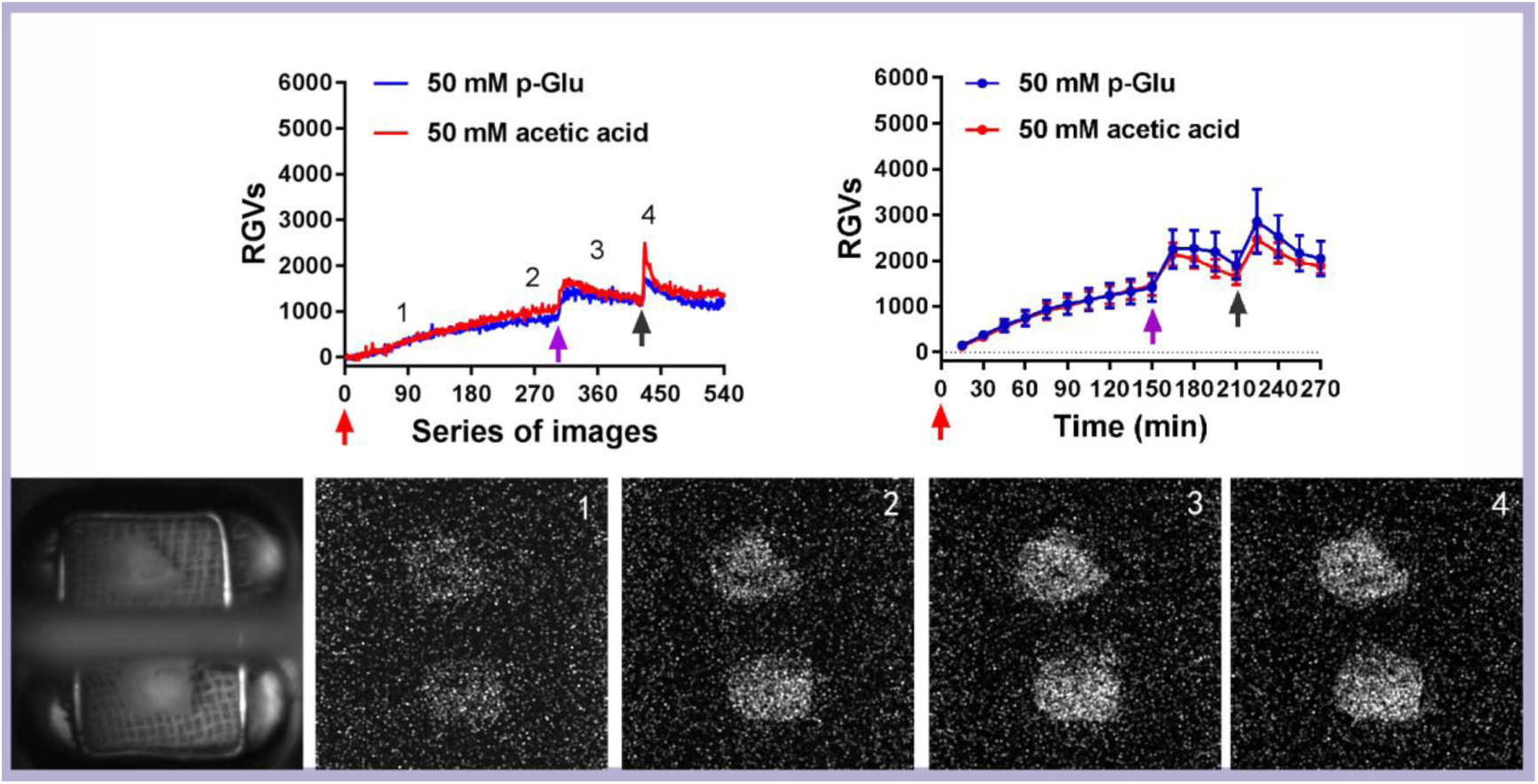
Acetic acid induced-biophotonic activity in mouse brain slices. Representative dynamic changes in the biophotonic activity demonstrated by RGVs are shown in left panel (30 s imaging time for each image). The sum of the time course of the average change in RGVs from 15 continuously processed original grey images is shown in right panel. Digit: 1-4 mark the 90, 270, 360, and 450 series of images, respectively, in the upper left panel, which corresponds to the four images in down panel. The red, pink and black arrows indicate the time points for application of 50 mM glutamate (upper image) and 50 mM acetic acid (0.3%, lower image) (0 min), washing (150 min) and reapplication (210 min), respectively. A representative regular image of two hemispherical slices (up and down) is shown on the left side of the lower panel. Acetic acid (50 mM) (red line, down image) also induced similar patterns of biophotonic activity compared to that of 50 mM p-Glu (blue line, upper image) (n=5). Data are shown as the mean±s. e. m. n=the number of slices from the same number of mice.

**Fig. S4.**
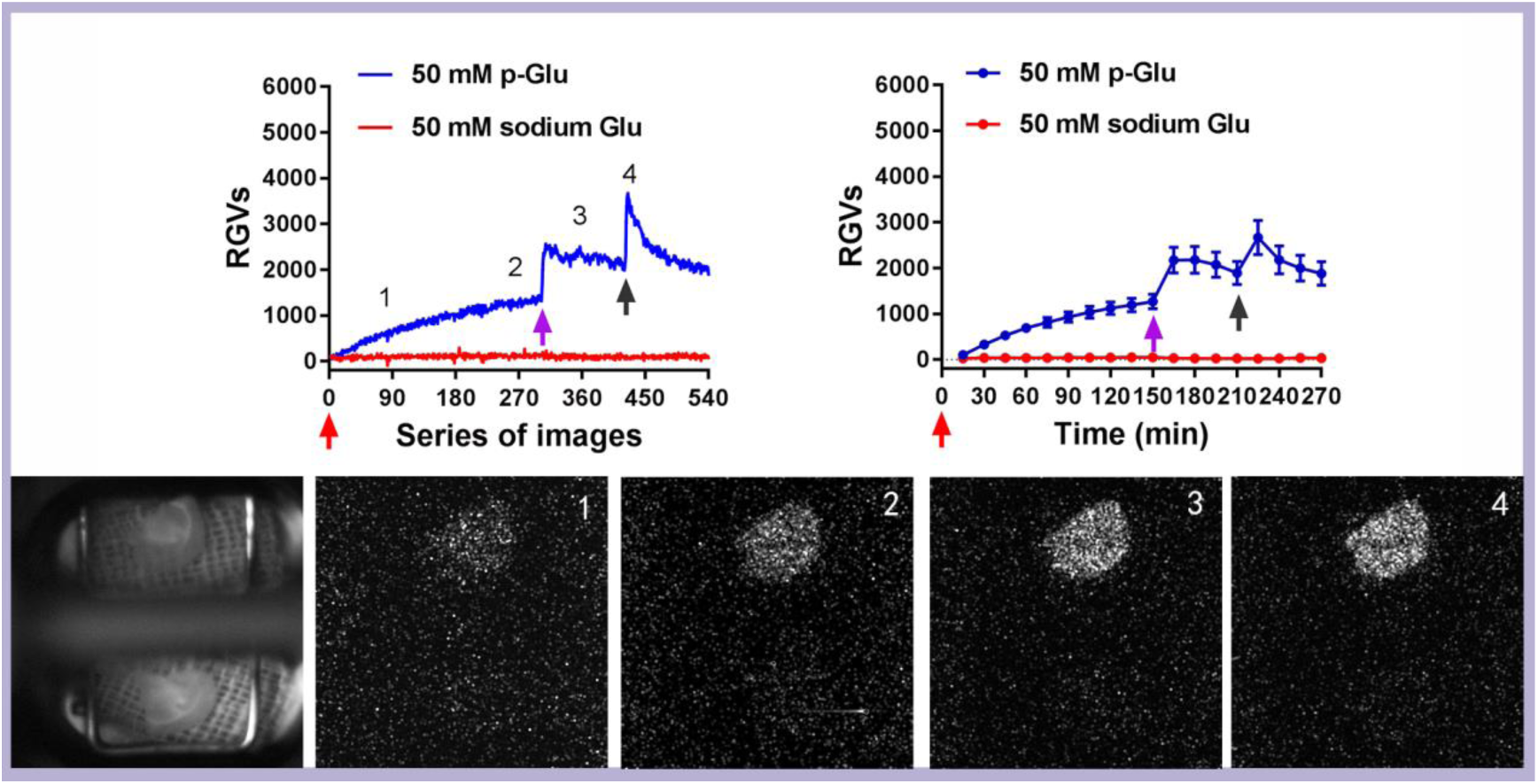
The effects of sodium glutamate on the biophotonic activity in mouse brain slices. Representative dynamic changes in the biophotonic activity demonstrated by RGVs are shown in the left panel (30 s imaging time for each image). The sum of the time course of the average change in RGVs from 30 continuously processed original grey images is shown in the right panel. Digit: 1-4 mark the 90, 270, 360, and 450 series of images, respectively, in upper left panel, which corresponds to the four images in down panel. The red, pink and black arrows indicate the time points for application of 50 mM glutamate (upper image) or 50 mM sodium glutamate (down image) (0 min), washing (150 min) and reapplication (210 min), respectively. A representative regular image of two hemispherical slices (up and down) is shown on the left side of the lower panel. Sodium glutamate (50 mM) (red line, down image) did not induce biophotonic activity, while 50 mM p-Glu (blue line, upper image) had typical effects (n=5). Data are shown as the mean±s. e. m. n=the number of slices from the same number of mice. Significant differences (* p<0.05 or ** p<0.01) are noted from 1-15 min (see also table S1).

**Fig. S5.**
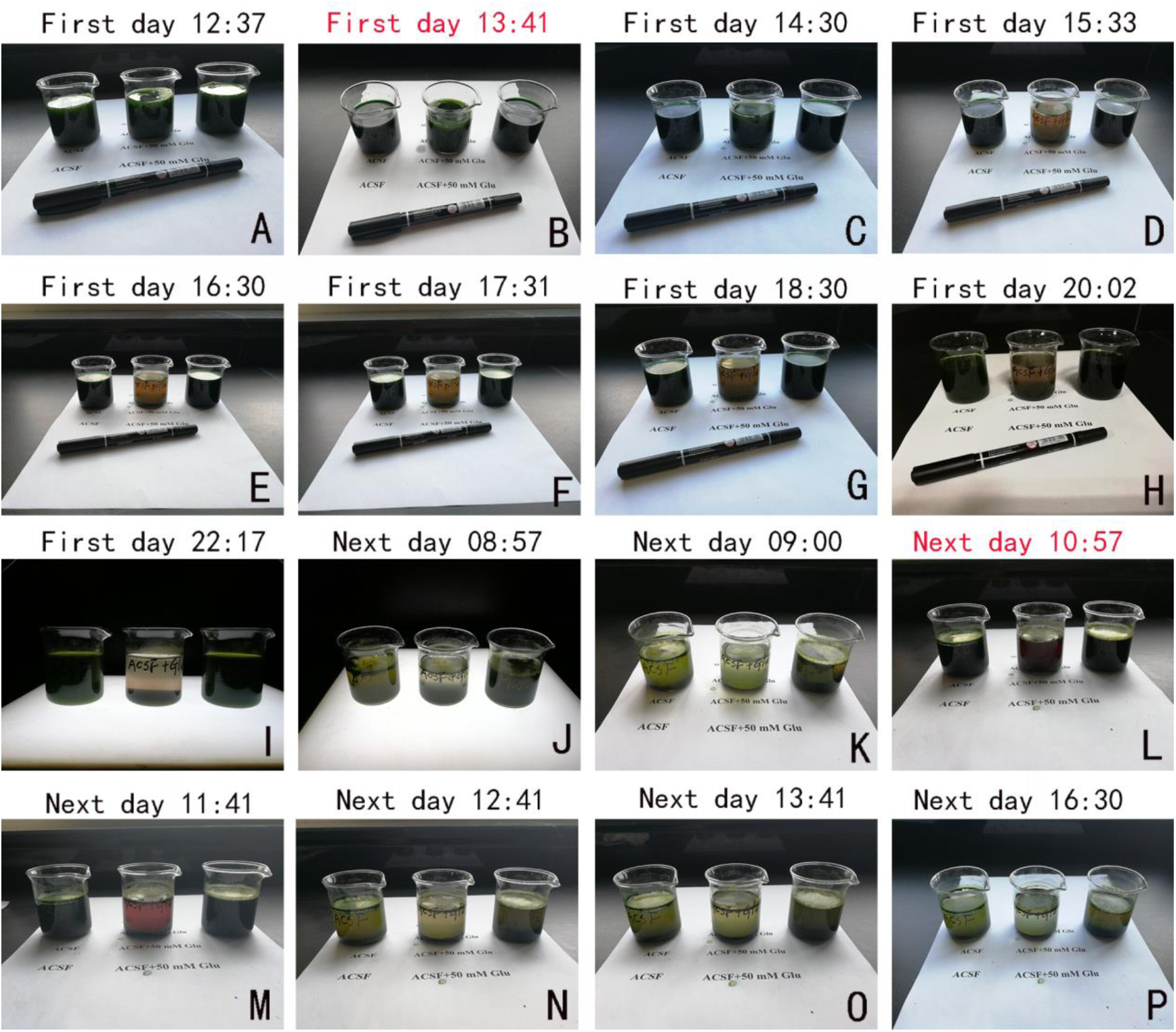
Determination of photochemical activity in ACSF and ACSF containing 50 mM glutamate. **(A-P)** Three reaction solutions (50 ml each) including ACSF, ACSF containing 50 mM p-Glu and reaction mixture [50 mM Tris-HCl buffer (pH 7.4), 0.4 mM sucrose, 10 mM NaC1] were placed into the left, middle and right bottles, respectively. **(A)** Freshly prepared chloroplast precipitate (1.5g) was added into the three reaction solutions (dark green). **(B)** One milliliter of 50 mM DCIP was added at 13:41 (highlights in red, dark blue). **(C-I)** The colour change from dark blue to purple-red-blue was observed for ACSF containing 50 mM p-Glu after incubation for approximately 2 h, but not for the ACSF and mixture solutions. **(J, K)** The colour change was obvious for three solutions after an overnight incubation for approximately 11 h under artificially controlled visible light irradiation. **(I)** One milliliter of 50 mM DCIP was added again at 10:57 the next day (dark blue). **(M-P)** The colour change was observed almost simultaneously in three reaction solutions after incubation for approximately 2 h, and the dark blue became colourless for both solutions containing ACSF.

**Table S1.** Statistical results for Fig.1-3 and Fig. S3, 4

| Time periods / Min | Comparison (two-tailed Student's t-test; p=) |  |  |  |  |  |  |  |  |
| --- | --- | --- | --- | --- | --- | --- | --- | --- | --- |
|  | Fig. 1B | Fig. 2C | Fig. 2D | Fig. 3C | Fig. 3C | Fig. 3C | Fig. 3D | Fig. S3 | Fig. S4 |
|  | P-Glu+ H89 vs P-Glu | p-Ic-Glu vs P-Glu | Ic-Glu vs P-Glu | Ic-Glu+ DC-ED vs Ic-Glu | Ic-Glu+ DC-ED vs P-Glu® | Ic-Glu+ DC-ED/ P-Glu® (%) | X vs Y | P-Glu vs AA | P-Glu vs S-Glu |
| 1-15 | 0.111061 | 0.046455 | 0.001041 | 0.017737 | 0.694645 | 85.61 | 0.265852 | 0.760842 | 0.019725 |
| 16-30 | 0.037493 | 0.074029 | 0.000209 | 0.000129 | 0.530209 | 118.25 | 0.335017 | 0.722619 | 0.001241 |
| 31-45 | 0.113286 | 0.082861 | 3.33E-05 | 0.000081 | 0.625955 | 111.19 | 0.346255 | 0.842564 | 0.000343 |
| 46-60 | 0.602259 | 0.107852 | 1.24E-05 | 0.000086 | 0.936222 | 101.64 | 0.323616 | 0.963451 | 0.000106 |
| 61-75 | 0.979144 | 0.097907 | 6.71E-06 | 0.000090 | 0.780611 | 94.65 | 0.319860 | 0.885215 | 0.000091 |
| 76-90 | 0.756283 | 0.070062 | 2.6E-06 | 0.000142 | 0.503719 | 87.48 | 0.174662 | 0.893319 | 0.000058 |
| 91-105 | 0.782199 | 0.044268 | 2.02E-06 | 0.000220 | 0.290115 | 80.86 | 0.272042 | 0.975382 | 0.000064 |
| 106-120 | 0.813402 | 0.018633 | 4.33E-07 | 0.000253 | 0.199645 | 77.49 | 0.317516 | 0.984990 | 0.000048 |
| 121-135 | 0.549230 | 0.008583 | 2.2E-07 | 0.000348 | 0.151917 | 75.43 | 0.279917 | 0.924295 | 0.000061 |
| 136-150 | 0.485377 | 0.005269 | 1.5E-07 | 0.000477 | 0.121066 | 73.96 | 0.344139 | 0.913217 | 0.000058 |
| 151-165 | 0.462758 | 0.001114 | 4.06E-08 | 0.000253 | 0.143753 | 75.92 | 0.154584 | 0.813503 | 0.000066 |
| 166-180 | 0.476307 | 0.003464 | 1.14E-07 | 0.000018 | 0.404149 | 84.07 | 0.104948 | 0.623459 | 0.000079 |
| 181-195 | 0.531320 | 0.003242 | 2.06E-07 | 0.000002 | 0.402031 | 83.25 | 0.236635 | 0.467502 | 0.000083 |
| 196-210 | 0.301633 | 0.002307 | 3.13E-07 | 0.000001 | 0.368346 | 82.00 | 0.213359 | 0.479112 | 0.000072 |
| 211-225 | 0.261704 | 0.000184 | 2.81E-07 | 0.000115 | 0.026793 | 59.00 | 0.220002 | 0.625199 | 0.000109 |
| 226-240 | 0.170134 | 0.000922 | 7.79E-07 | 0.000112 | 0.074292 | 65.85 | 0.258876 | 0.513527 | 0.000105 |
| 241-255 | 0.431832 | 0.001238 | 5.59E-07 | 0.000085 | 0.102170 | 69.56 | 0.301080 | 0.655236 | 0.000115 |
| 256-270 | 0.276195 | 0.001004 | 4.98E-07 | 0.000088 | 0.087547 | 70.10 | 0.236032 | 0.720233 | 0.000094 |
| 271-285 | 0.002918 |  |  |  |  |  |  |  |  |
| 286-300 | 0.000944 |  |  |  |  |  |  |  |  |
| 301-315 | 0.006127 |  |  |  |  |  |  |  |  |
| 316-330 | 0.009609 |  |  |  |  |  |  |  |  |
| 331-345 | 0.008212 |  |  |  |  |  |  |  |  |
| 346-360 | 0.004747 |  |  |  |  |  |  |  |  |
| 361-375 | 0.003466 |  |  |  |  |  |  |  |  |
| n | (6, 6) | (6, 6) | (6, 6) | (8, 8) | (8, 37) | (8, 37) | (5, 5) | (5, 5) | (5, 5) |
| P-Glu: purchased glutamate; Ic-Glu: incompetent glutamate (P-Glu+4whole brain slices); p-Ic-Glu: hypofunctional or partial incompetent glutamate (P-Glu+2 whole brain slices); P-Glu®: the reference data induced by 50 mM p-Glu; ACSF: artificial cerebral spinal fluid; DC-ED: direct current electrical discharges; X: ACSF+4 whole brain slices+DC-ED; Y: ACSF+4 whole brain slices; AA: acetic acid; S-Glu: sodium glutamate; n: the number of slices from the same number of mice; Ex-Fig: Extended Data-Fig; Highlights: blue<0.05, red<0.01. |  |  |  |  |  |  |  |  |  |

**Table S2.** Statistical results for Fig. 4

| Time periods / Min | Comparison (two-tailed Student's t-test; p=) |  |  |  |  |  |  |  |  |
| --- | --- | --- | --- | --- | --- | --- | --- | --- | --- |
|  | Fig. 4A | Fig. 4A | Fig. 4A | Fig. 4B | Fig. 4B | Fig. 4B | Fig. 4C | Fig. 4C | Fig. 4C |
|  | Rc-Glu | Rc-Glu | Rc-Glu/ | A | A | B | C | C | D |
|  | vs | vs | P-Glu® | vs | vs | vs | vs | vs | vs |
|  | Ic-Glu | P-Glu® | (%) | B | P-Glu® | P-Glu® | D | P-Glu® | P-Glu® |
| 1-15 | 0.000446 | 0.000003 | 352.89 | 0.001545 | 0.802369 | 0.013355 | 0.001238 | 0.001759 | 0.114614 |
| 16-30 | 0.000573 | 0.000000 | 370.70 | 0.000523 | 0.917060 | 0.004007 | 0.000664 | 0.000406 | 0.045860 |
| 31-45 | 0.000181 | 0.000000 | 300.96 | 0.000569 | 0.777067 | 0.000768 | 0.000508 | 0.001131 | 0.014505 |
| 46-60 | 0.000063 | 0.000000 | 260.51 | 0.000596 | 0.906231 | 0.000237 | 0.000804 | 0.007962 | 0.008987 |
| 61-75 | 0.000049 | 0.000002 | 233.30 | 0.00039 | 0.956692 | 0.000097 | 0.001625 | 0.054312 | 0.007349 |
| 76-90 | 0.000026 | 0.000007 | 218.09 | 0.000366 | 0.842642 | 0.000092 | 0.003295 | 0.152078 | 0.013953 |
| 91-105 | 0.000017 | 0.000012 | 208.41 | 0.000373 | 0.699655 | 0.000066 | 0.008907 | 0.291834 | 0.022430 |
| 106-120 | 0.000020 | 0.000023 | 200.72 | 0.000196 | 0.701848 | 0.000041 | 0.017878 | 0.423368 | 0.038372 |
| 121-135 | 0.000023 | 0.000030 | 196.01 | 0.000141 | 0.659422 | 0.000028 | 0.030618 | 0.511117 | 0.058459 |
| 136-150 | 0.000017 | 0.000046 | 190.59 | 4.48E-05 | 0.667130 | 0.000019 | 0.051075 | 0.628595 | 0.071337 |
| 151-165 | 0.000016 | 0.000302 | 177.64 | 0.000211 | 0.163231 | 0.000019 | 0.005703 | 0.925506 | 0.000948 |
| 166-180 | 0.000010 | 0.000097 | 203.21 | 0.00113 | 0.146955 | 0.000295 | 0.000913 | 0.816077 | 0.001961 |
| 181-195 | 0.000005 | 0.000353 | 197.93 | 0.001844 | 0.146365 | 0.000601 | 0.000738 | 0.969843 | 0.002460 |
| 196-210 | 0.000003 | 0.001465 | 185.42 | 0.002024 | 0.143301 | 0.000615 | 0.000786 | 0.748297 | 0.002075 |
| 211-225 | 0.000012 | 0.010940 | 160.25 | 0.001588 | 0.013288 | 0.000177 | 0.000661 | 0.700335 | 0.000285 |
| 226-240 | 0.000020 | 0.000672 | 187.65 | 0.001014 | 0.027368 | 0.000273 | 0.000179 | 0.613401 | 0.000918 |
| 241-255 | 0.000008 | 0.000307 | 190.89 | 0.000758 | 0.038213 | 0.000189 | 0.000165 | 0.477049 | 0.001558 |
| 256-270 | 0.000005 | 0.000270 | 185.54 | 0.000862 | 0.035565 | 0.000077 | 0.000167 | 0.522276 | 0.001284 |
| n | (5, 5) | (5, 37) | (5, 37) | (4, 4) | (4, 37) | (4, 37) | (5, 5) | (5, 37) | (5, 37) |
| Ic-Glu: incompetent glutamate; Rc-Glu: recompetent glutamate (Ic-Glu+Chloroplasts); P-Glu: purchased glutamate; P-Glu®: the reference data induced by 50 mM p-Glu; ACSF: artificial cerebral spinal fluid; A: ACSF+4 whole brain slices+chloroplasts; B: ACSF+4 whole brain slices; C: ACSF+chloroplasts; D: ACSF+chloroplasts+adjusting pH to 7.4; n: the number of slices from the same number of mice; Highlights: *blue<0.05, **red<0.01. |  |  |  |  |  |  |  |  |  |

## References

1. Lee EJD, Williams KM (1990) Chirality. Clinical pharmacokinetic and pharmacodynamic considerations. Clin Pharmacokinet. 18:339–345.

2. Sell CS (2006) On the unpredictability of odor. Angew. Chem. 45:6254–6261.

3. Brookes JC (2016) Quantum effects in biology: golden rule in enzymes, olfaction, photosynthesis and magnetodetection. Proc Math Phys Eng Sci. 473:20160822.

4. Franco MI, Turin L, Mershin A, Skoulakis EMC (2011) Molecular vibration-sensing component in Drosophila melanogaster olfaction. Proc. Natl Acad Sci. USA. 108:3797–3802.

5. Michaeli K, Kantor-Uriel N, Naaman R, Waldeck DH (2016) The electron's spin and molecular chirality - how are they related and how do they affect life processes?. Chem Soc Rev. 45:6478–6487.

6. Wolynes PG (2009). Some quantum weirdness in physiology. Proc Natl Acad Sci U S A. 106:17247–17248.

7. Lambert N. et al. (2013) Quantum biology. Nature Physics 9:10–18.

8. Tang R, Dai J (2014) Spatiotemporal imaging of glutamate-induced biophotonic activities and transmission in neural circuits. PloS one 9:e85643.

9. Tang R, Dai J (2014) Biophoton signal transmission and processing in the brain. J Photochem Photobiol B. 5:139:71–75.

10. Salari V, Valian H, Bassereh H, Bókkon I, Barkhordari A (2015) Ultraweak photon emission in the brain. J Integr Neurosci. 14:419–429.

11. Kumar S, Boone K, Tuszyński J, Barclay P, Simon C (2016) Possible existence of optical communication channels in the brain. Sci Rep. 6:36508.

12. Chai W. et al. (2018) Biophotonic activity and transmission mediated by mutual actions of neurotransmitters are involved in the origin and altered states of consciousness. Neurosci Bull. 34:534–538.

13. Wang Z, Wang N, Li Z, Xiao F, Dai J (2016) Human high intelligence is involved in spectral redshift of biophotonic activities in the brain. Proc. Nat. Acad Sci U S A 113:8753–8758.

14. Tamminga CA (1998) Schizophrenia and glutamatergic transmission. Crit Rev Neurobiol. 12: 21–36.

15. Olney JW, Newcomer JW, Farber NB (1999) NMDA receptor hypofunction model of schizophrenia. J Psychiatr Res. 33:523–533.

16. Kantrowitz J, Javitt DC (2012) Glutamatergic transmission in schizophrenia: From basic research to clinical practice. Curr Opin Psychiatry 25:96–102.

17. Gilmour G. et al. (2012) NMDA receptors, cognition and schizophrenia--testing the validity of the NMDA receptor hypofunction hypothesis. Neuropharmacology 62:1401–1412.

18. Snyder MA, Adelman AE, Gao WJ (2013) Gestational methylazoxymethanol exposure leads to NMDAR dysfunction in hippocampus during early development and lasting deficits in learning. Neuropsychopharmacology 38:328–340.

19. Catts VS, Lai YL, Weickert CS, Weickert TW, Catts SVA (2016) Quantitative review of the postmortem evidence for decreased cortical N-methyl-D-aspartate receptor expression levels in schizophrenia: How can we link molecular abnormalities to mismatch negativity deficits? Biol Psycho!. 116:57–67.

20. Kosten L. et al. (2016) Multiprobe molecular imaging of an NMDA receptor hypofunction rat model for glutamatergic dysfunction. Psychiatry Res Neuroimaging 248:1–11.

21. Rappeneau V, Blaker A, Petro JR, Yamamoto BK, Shimamoto A (2016) Disruption of the glutamate-glutamine cycle involving astrocytes in an animal model of depression for males and females. Front Behav Neurosci. 10:e23l.

22. Bardoni R, Torsney C, Tong CK, Prandini M, MacDermott AB (2004) Presynaptic NMDA receptors modulate glutamate release from primary sensory neurons in rat spinal cord dorsal horn. J. Neurosci. 24:2774–2781.

23. Engel GS. et al. (2007) Evidence for wavelike energy transfer through quantum coherence in photosynthetic systems. Nature 446:782–786.

24. Panitchayangkoon G. et al. (2010) Long-lived quantum coherence in photosynthetic complexes at physiological temperature. Proc. Nat. Acad. Sci U S A 107:12766–12770.

25. Tempelaar R, Jansen TLC, Knoester J (2014) Vibrational beatings conceal evidence of electronic coherence in the FMO light-harvesting complex. J. Phys. Chem. B. 118:12865–12872.

26. Monahan DM, Whaley-Mayda L, Ishizaki A, Fleming GR (2015) Influence of weak vibrational-electronic couplings on 2D electronic spectra and inter-site coherence in weakly coupled photosynthetic complexes. J. Chem. Phys. 143:065101.

27. Kaniuga Zl, Sochanowicz B, Zabek J, Krystyniak K (1978) Photosynthetic apparatus in chilling-sensitive plants: I. Reactivation of hill reaction activity inhibited on the cold and dark storage of detached leaves and intact plants. Planta 140:121–128.

28. Yin ZH, Dietz KJ, Heber U (1990) Light-dependent pH changes in leaves of C3 plants: III. Effect of inhibitors of photosynthesis and of the developmental state of the photosynthetic apparatus on cytosolic and vacuolar pH changes. Planta 182:262–269.

29. Shikanai T, Yamamoto H (2017) Contribution of cyclic and pseudo-cyclic electron transport to the formation of proton motive force in chloroplasts. Mol Plant. 10:20–29.

30. Davis GAet al. (2016) Limitations to photosynthesis by proton motive force-induced photosystem II photodamage. Elife 5:e16921.

31. Gray HB, Winkler JR (2003) Electron tunneling through proteins. Q Rev Biophys. 36:341–372.

32. Nagel ZD, Klinman JP (2006) Tunneling and dynamics in enzymatic hydride transfer. Chem Rev. 106:3095–3118.

33. Halpin A et al. (2014) Two-dimensional spectroscopy of a molecular dimer unveils the effects of vibronic coupling on exciton coherences. Nature Chemistry 6:196–201.

34. Danbolt NC (2001) Glutamate uptake. Prog. Neurobiol. 65:101–105.

35. Jiang J, Amara SG (2011) New views of glutamate transporter structure and function: advances and challenges. Neuropharmacology 60:172–181.

36. Eitan R, Lerer B (2006) Nonpharmacological, somatic treatments of depression: electroconvulsive therapy and novel brain stimulation modalities. Dialogues Clin Neurosci. 8:241–258.

37. Yrondi A. et al. (2018) Electroconvulsive therapy, depression, the immune system and inflammation: A systematic review. Brain Stimul. 11:29–51.

## References

1. Tang R, Dai J (2014) Spatiotemporal imaging of glutamate-induced biophotonic activities and transmission in neural circuits. PloS one 9:e85643.

2. Wang Z, Wang N, Li Z, Xiao F, Dai J (2016) Human high intelligence is involved in spectral redshift of biophotonic activities in the brain. Proc. Nat. Acad. Sci U S A 113:8753–8758.

3. Kaniuga Zl, Sochanowicz B, Zabek J, Krystyniak K (1978) Photosynthetic apparatus in chilling-sensitive plants: I. Reactivation of hill reaction activity inhibited on the cold and dark storage of detached leaves and intact plants. Planta 140:121–128.

